# Inferring delays in partially observed gene regulatory networks

**DOI:** 10.1101/2022.11.27.518074

**Authors:** Hyukpyo Hong, Mark Jayson Cortez, Yu-Yu Cheng, Hang Joon Kim, Boseung Choi, Krešimir Josić, Jae Kyoung Kim

## Abstract

**Motivation:** Cell function is regulated by gene regulatory networks (GRNs) defined by protein-mediated interaction between constituent genes. Despite advances in experimental techniques, we can still measure only a fraction of the processes that govern GRN dynamics. To infer the properties of GRNs using partial observation, unobserved sequential processes can be replaced with distributed time delays, yielding non-Markovian models. Inference methods based on the resulting model suffer from the curse of dimensionality.

**Results:** We develop a simulation-based Bayesian MCMC method for the efficient and accurate inference of GRN parameters when only some of their products are observed. We illustrate our approach using a two-step activation model: An activation signal leads to the accumulation of an unobserved regulatory protein, which triggers the expression of observed fluorescent proteins. With prior information about observed fluorescent protein synthesis, our method successfully infers the dynamics of the unobserved regulatory protein. We can estimate the delay and kinetic parameters characterizing target regulation including transcription, translation, and target searching of an unobserved protein from experimental measurements of the products of its target gene. Our method is scalable and can be used to analyze non-Markovian models with hidden components.

**Availability:** Accompanying code in R is available at https://github.com/Mathbiomed/SimMCMC.

**Supplementary information:** Supplementary data are available at *Bioinformatics* online.

## 1 Introduction

Advances in microscopy allow us to observe the dynamics of gene regulatory networks (GRNs) in unprecedented detail. Novel statistical techniques have helped interpret the resulting wealth of data. However, even the best experimental methods can provide observations of only a fraction of the components constituting a GRN, and thus offer only partial information about the dynamics of gene circuits. Statistical methods thus need to take into account the effect of unobserved processes to correctly interpret the data, and accurately characterize genetic circuits and their dynamics.

Recently, inference methods have been proposed based on the assumption that the unobserved processes are sequential, and thus can be modeled by introducing a delay (Jiang *et al*., 2021; Heron *et al*., 2007; Calderazzo et *al*., 2018; Choi *et al*., 2020; Cortez *et al*., 2022; Barrio *et al*., 2013; Leier *et al*., 2014; Gomez *et al*., 2016; Kim *et al*., 2022). The resulting models are non-Markovian, as system dynamics depends not only on the present, but also past states. This model drastically reduces the number of parameters and reactions that need to be inferred. However, this approach can only account for effects of sequential processes such as transcription and translation, which are modeled as delays in interactions between the genes in the network (Fig. 1). Thus, it is unclear how to analyze data and perform inference when the products of some genes in the circuit are unobserved. For example, in a network of two genes *x* and y, where protein X regulates the expression of gene *y*, present methods can be applied when counts of both proteins X and Y are known. Often it is impossible to observe the products of both genes concurrently. Is it possible to characterize the dynamics of X, if only Y is observed?

**Fig. 1.**
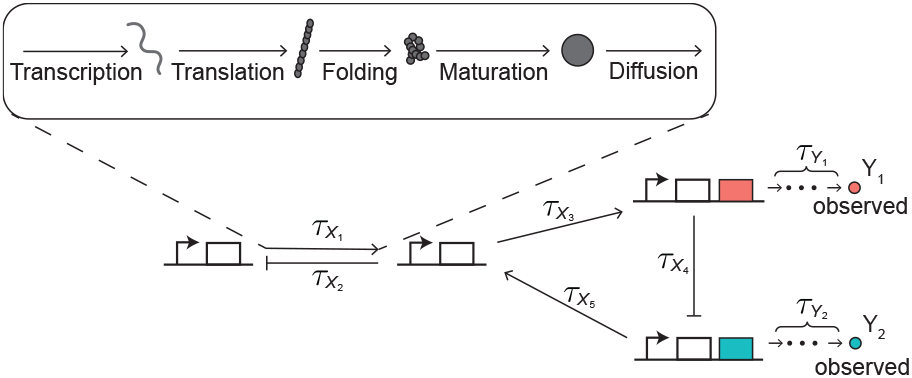
Gene regulatory networks consist of genes whose interactions can be described with distributed time delays. In such networks, we can often observe the product (i.e., protein) of only some genes by using fluorescence microscopy (Y_1_ and Y_2_). Protein synthesis consists of multiple steps (transcription, translation, folding, maturation), and its duration can be described using a distributed time delays (*τ*_Y_i__). Further delays, *τ*_X_i__, can also be used to describe interactions between unobserved and observed genes. Such delays take into account protein synthesis, 3D diffusion inside a cell, and sliding along a strand of DNA to find a promoter region.

Currently available Bayesian Markov chain Monte Carlo (MCMC) methods for inference of non-Markovian systems are not well suited to address this question. While these methods are applicable even to systems with high intrinsic noise (i.e., low copy numbers of molecules) (Choi *et al*., 2020) and cell-to-cell heterogeneity (Cortez *et al*., 2022), they rely on the assumption that all proteins in a GRN are observed. When some protein counts are unobserved, extending such methods directly requires that we characterize reactions involving the unobserved proteins at each measurement time. The high dimensionality of the resulting system makes inference challenging, if not impossible.

Here, we present a simulation-based Bayesian method for the inference of kinetic and delay parameters of a GRN when only the products of some of the genes in the network are observed. The approach is applicable generally even if only the most downstream genes, i.e., the final outputs, of the network are observed. We illustrate the method using a two-step activation model, where an initial signal activates gene *x* whose product protein X is unobserved. Protein X triggers the expression of gene *y* and the production of an observed protein Y. By performing an identifiability analysis, we characterize what information about protein Y dynamics is needed to obtain accurate and precise information about the unobserved protein X. We find that the dilution rate and synthesis time delay of Y need to be known in order to infer the kinetic and time delay parameters characterizing the dynamics of X. We apply this approach to a plasmid-borne two-step activation circuit in *Escherichia coli (E. coli)* where unobserved AraC protein triggers the expression of yellow fluorescent protein (YFP). When the dilution rate and the time delay for the synthesis of the observed protein YFP are separately measured, we are able to infer the time delay for target regulation by the unobserved AraC protein (i.e., the delay due to transcription, translation, and target searching). This finding can play a critical role in synthetic circuit design because the AraC protein, whose kinetics has yet to be fully characterized, is a widely used transcriptional activator in synthetic biology. Our study also illustrates how information from unobserved proteins can be inferred from the dynamics of observed proteins in GRNs. Our approach is scalable and provides a tool for characterizing non-Markovian systems from partial observations.

## 2 Materials and methods

### 2.1 Derivation of a likelihood function of kinetic and delay parameters

We first derive a likelihood function to construct a Bayesian inference method for estimating kinetic and delay parameters (Choi *et al*., 2017, 2020; Hong *et al*., 2022; Cortez *et al*., 2022). Consider a biochemical reaction network with *u* species *Z*_1_,…,*Z_u_* and *v* reactions *R*_1_,…,*R_v_*. Reaction *R_k_* can be represented as

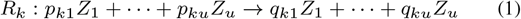

where *p_kj_* and *q_kj_* are the stoichiometric coefficients of species *Z_j_*. For each reaction *R_k_*, the reaction initiation rate, *h_k_*(*z*(*t*), *θ_k_*), is a function of the current state *z*(*t*) = (*z*_1_(*t*),…, *z_u_*(*t*)) and the associated kinetic parameters *θ_k_*, where *z_j_*(*t*) is the number of species *Z_j_* at time *t*. We assume that each reaction takes a random time to complete. Therefore, after a reaction is initiated, the system state changes only after a random delay. This delay follows a distribution *η_k_* fully determined by a vector of parameters, Δ_*k*_. For example, in a GRN, when the synthesis of a transcriptional activator protein is initiated, a functional protein is produced after a sequence of steps including transcription, translation, and maturation (Golding *et al*., 2005; Kærn *et al*., 2005). Each of these steps takes time, and only after all steps are completed can the functional protein diffuse to its target binding site (Cheng *et al*., 2017; Elf *et al*., 2007; Hammar *et al*., 2012). Thus, the number of functional activator proteins increases only when all intermediate steps are completed. If a reaction is not delayed, then the associated delay distribution is the Dirac delta measure at time 0.

Schlicht and Winkler, 2008 have proven that a reaction *completion* propensity, *f_k_* (*t*, **z**_c_, *θ_k_*, Δ_*k*_), describes the effective reaction rate of *R_k_* at time *t*. This propensity depends on **z**_c_, the *complete* trajectory of all species from time 0 to the maximum measurement time *T*. The reaction completion propensity is a function of the reaction initiation propensity, *h*, and the delay distribution, *η*, and is given by:

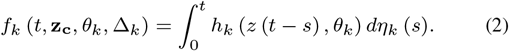

Intuitively, the completion propensity is an average of past reaction initiation propensities weighted by the probability that they have occurred a given time in the past.

We can define the likelihood of the kinetic and delay parameters, *θ* and **Δ**, respectively, for the given trajectory **z**_c_ (Gupta and Rawlings, 2014):

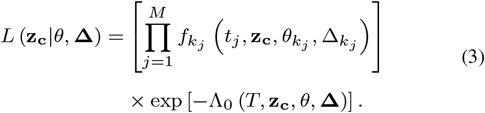

Here the pair (*t_j_*, *k_j_*) for *j* = 1,…, *M* denotes the completion time and type of a reaction that completes within the time interval (0, *T*]. In addition,

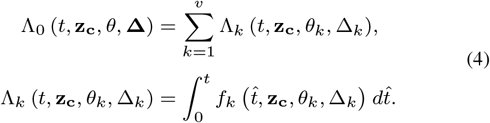

This is analogous to the likelihood provided by Boys *et al*. (2008) for a system without delays.

The trajectories of biochemical species can be measured experimentally only at discrete time points. Complete reaction histories are thus unknown. If we measure the trajectories at discrete time points *t* = 0,…, *T*, and denote these measurements by **z**, then according to Choi *et al*. (2020); Cortez *et al*. (2022), an approximate likelihood function, 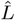, is given by

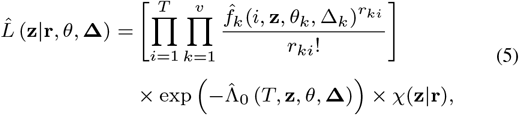

where **r** = (**r**_1_,…,**r_*v*_**) is a vector of reaction counts completing within each of the time intervals, i.e., **r**_*k*_ = (*r*_*k*1_,…, *r_kT_*) and *r_ki_* is the count of reaction of type *k* that completes within the time interval (*i* – 1, *i*]. Note that *χ*(**z**|**r**) is the indicator function that is one if the trajectory of species matches the reaction counts and zero otherwise (see Supplementary Methods for details). In Eq. (5), 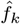 is an approximate reaction completion propensity computed by linearly interpolating the exact completion propensity (Eq. (2)) using:

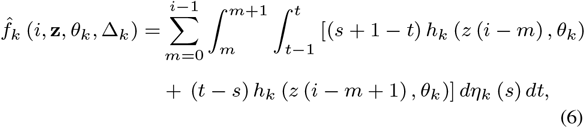

and 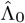 is defined analogously to Λ_0_ with 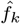 replacing *f_k_* in Eq. (4).

Since the approximate likelihood given in Eq. (5) takes into account the number of reactions completed between discrete time points, it corresponds to the *τ*-leaping approach (Gillespie, 2001). The exact likelihood in Eq. (3) corresponds to the exact stochastic simulation algorithm (SSA) for a system with delays (Cai, 2007).

### 2.2 Derivation of a likelihood function given noisy observations of a subset of species

In GRNs, we cannot measure the activity of all components directly. However, we can measure the activity of fluorescent reporter proteins (Fig. 1). These measurements are often contaminated by observational noise, which we assume is characterized by the vector of noise parameters σ. We derive a likelihood function for these noisy observations, assuming that only some protein counts are observed.

Let **x** and **y** be the trajectories of the unobserved and observed species, respectively, in a biochemical reaction network with delays so that **z** = (**x**, **y**). We let **y**_obs_ be the vector of noisy observations of species **y** at discrete time points *t* = 0,…, *T*. We assume that **y**_obs_ is obtained by adding i.i.d. noises from a Gaussian distribution to each observation in **y**. The joint likelihood function of the unknown kinetic and delay parameters and reaction counts is then given by

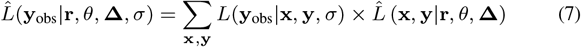

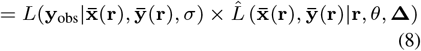

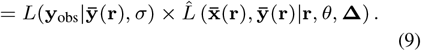

The sum over **y** in Eq. (7) can be reduced to a single term in Eq. (8) because the vector of reaction counts, **r**, uniquely determines the trajectory in the indicator function in Eq. (5), *χ*(**x**, **y**|**r**). Thus, the other terms in the sum vanish, and we denote the trajectories of unobserved and observed species without noise matching the vector of reaction counts by 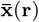 and **ȳ**(**r**), respectively. Furthermore, Eq. (8) simplifies to Eq. (9) as the noisy observations depend only on the trajectory of the observed species, **y**(**r**), and are thus conditionally independent of **x**(**r**). On the right-hand side of Eq. (9), the first factor is the likelihood function of the noisy observation of the given trajectory at discrete time points, **y**_obs_, while the second factor is the approximate likelihood function given in Eq. (5). Based on this approximate joint likelihood, we developed a Bayesian MCMC algorithm for a delayed reaction system with noisy measurements of the observed components.

### 2.3 Simulation-based MCMC for discrete noisy observation from a gene regulatory network with time delays

Sampling from the conditional posterior distribution of the parameters characterizing the unobserved processes requires protein counts as input. However, because we cannot measure all protein counts directly, we have to generate samples of the unobserved protein counts as well. As we explain below, the random walk approach used previously for this purpose suffers from the curse of dimensionality (Boys *et al*., 2008; Choi *et al*., 2020; Cortez *et al*., 2022). To circumvent this problem, we use stochastic simulations to generate samples of the unobserved protein counts. We describe the general idea behind our approach in this section. We provide details and the example of the two-activation model in the Supplementary Methods.

Using Bayes’ theorem, we can obtain the joint posterior distribution of the model parameters, (*θ*, **Δ**), and the number of reactions, **r**, given the noisy measurements of the observed species, **y**_obs_, by multiplying the priors and the joint likelihood of the unknowns (Eq. (9)):

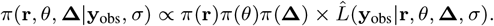

When we generate samples from this joint posterior distribution, the dimension of the sampling distribution increases with the number of parameters and measurements. To generate samples in high dimensions, we exploit the Gibbs sampling approach and decompose each sampling step in the high dimensional space into separate low dimensional sampling steps. We use the Metropolis-Hastings (MH) -within-Gibbs sampler method to sample from each conditional posterior.

Although we divided each high dimensional sampling step into iterative low dimensional sampling steps, there are other practical challenges to implementing the block updating method we have used previously (Boys *et al*., 2008; Choi *et al*., 2020; Cortez et al., 2022). First, because the block updating method is based on the MH algorithm, we need to tune too many hyperparameters (i.e., the variances of proposal distributions). Second, even if we tuned all proposal distributions, the MCMC algorithm converges slowly because all reaction counts are updated independently, while the reaction counts on subsequent time intervals are strongly correlated. Thus, using the random walk chain for each reaction count can significantly reduce the acceptance probability of proposed samples, leading to slow convergence of the proposed MCMC algorithm.

To address these problems, we utilize an algorithm that generates proposal reaction counts based on simulations of the biochemical reaction network in Eq. (1) (Wilkinson, 2018). Simulating the model directly obviates the need for parameter tuning and captures correlations between the reaction counts on subsequent time intervals. For simulations we chose τ-leaping (Gillespie, 2001) because it corresponds to the approximate likelihood in Eq. (9) and it is computationally more efficient than the exact delayed SSA (Cai, 2007).

The resulting MCMC procedure can be described as follows (Fig. 2), with the superscript (*j*) denoting samples at the *j*-th MCMC iteration step:

1. Initialize the kinetic and delay parameters (*θ*^(0)^, **Δ**^(0)^) and reaction counts **r**^(0)^.
2. By performing the stochastic simulation for given (*θ*^(*j*)^, **Δ**^(*j*)^), propose new trajectories **x**\* and **y**\* and the underlying reaction counts **r**\*.
3. Accept the proposed reaction counts and corresponding trajectories (**r**\*, **x**\*, **y**\*) based on the MH acceptance probability of *ρ*(**r**\*, **r**^(*j*)^) where

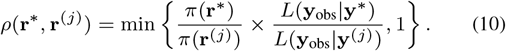
4. Sample the kinetic and delay parameters (θ^(*j*+1)^, **Δ**^(*j*+1)^) from their full conditional posterior distribution for the given reaction counts **r**^(*j*+1)^ using an MH-within-Gibbs sampling approach.
5. Repeat Steps 2–4 until a convergence criterion is met.

**Fig. 2.**
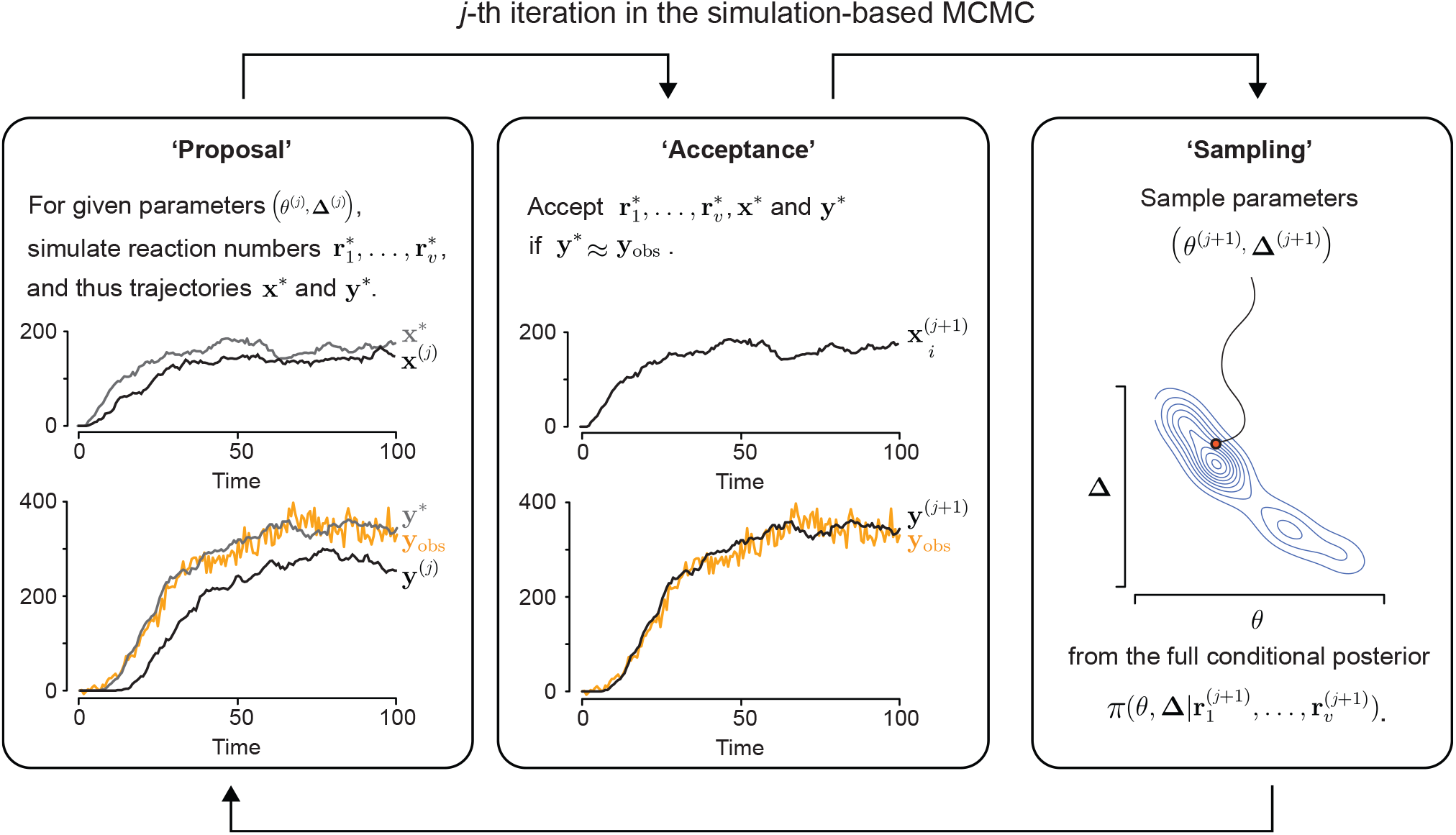
An illustration of the simulation-based MCMC method to estimate the posterior distribution of the kinetic (*θ*) and delay parameters (**Δ**) of a GRN with unobserved, *X*, and observed, *Y*, components. (**Proposal**) At the *j*-th MCMC iteration, for given parameter samples, *θ*^(*j*)^ and **Δ**^(*j*)^, we simulate a stochastic model with time delays and propose candidates for the reaction counts in the model (i.e., 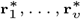), which uniquely determine the trajectories for X and Y (i.e., **x**\* and **y**\*). (**Acceptance**) The proposed reaction counts and trajectories are more likely to be accepted if **y**\* is closer to the observation **y**_obs_ than the previous sample **y**^(*j*)^ (see text and Materials and methods for details). If the proposed reaction counts and trajectories are not accepted, those from the previous iteration are kept as the current samples. The updated reaction counts and trajectories are referred to as 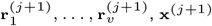, and **y**^(*j*+1)^. (**Sampling**) For given updated reaction counts, we sample the kinetic and delay parameters from their full conditional posterior distribution 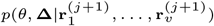 using MH-within-Gibbs sampling approach. These steps are repeated until a convergence criterion is met.

The acceptance probability in Eq. (10) can be viewed as the conventional MH acceptance probability when a sample from a proposal distribution is replaced with a sample generated by a stochastic simulation. We provide the derivation of the acceptance probability, *ρ*(**r**\*, **r**^(*j*)^), and the explicit form of conditional posterior distributions of the kinetic and delay parameters in Step 4, *π*(*θ*, **Δ**|**r**^(*j*+1)^), in the Supplementary Methods.

## 3 Results

### 3.1 Kinetic and delay parameters can be accurately estimated as long as unidentifiable parameters are known

We first applied our inference algorithm to synthetic data obtained from simulations of a two-step activation model (Fig. 3a) to test if our method can accurately estimate the kinetic and delay parameters of a GRN. In the generative model, the production of the unobserved protein X is initiated at rate of *A_X_* and the protein activates the production of the observed protein Y. Transcription of Y is initiated at a rate that is modeled by a Michaelis-Menten function, after a regulation delay *τ_X_*. Protein Y takes a random time to mature, and hence each molecule becomes observable after a time delay, *τ_Y_*. Proteins X and Y are diluted at the same rate *B* due to cell growth. When trying to infer all parameters in this model, we encountered identifiability issues. However, we show that we can obtain accurate and precise parameters estimates when some of the unidentifiable parameters are known.

**Fig. 3.**
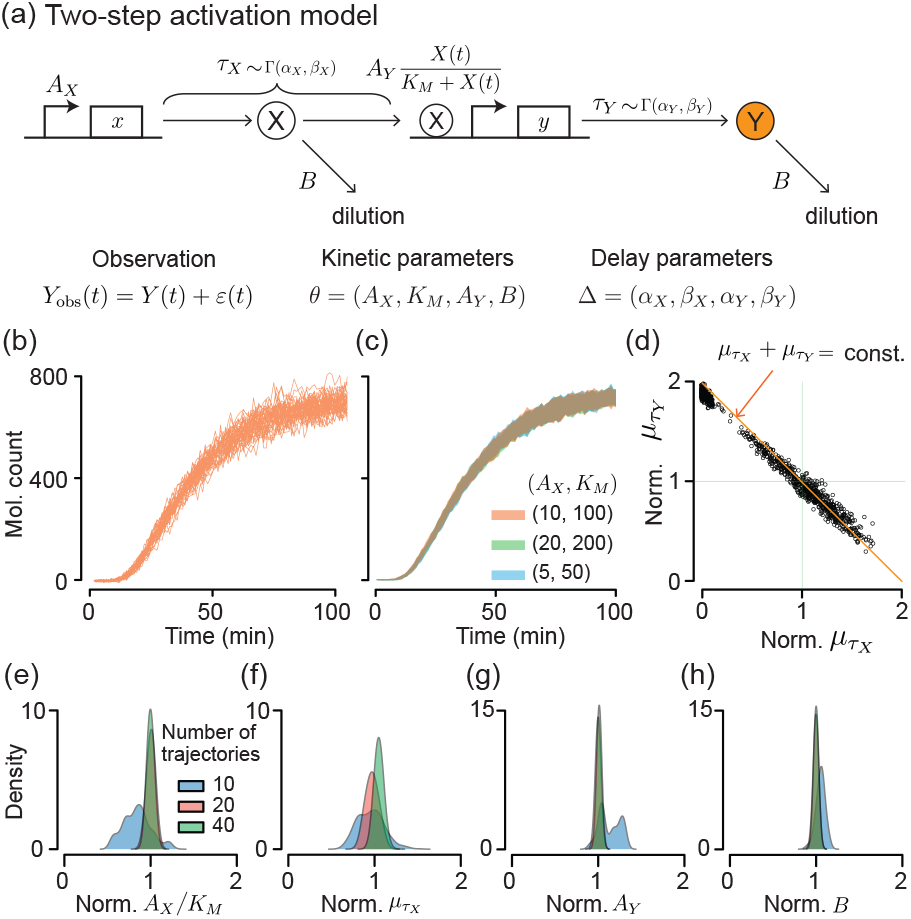
Estimation of the kinetic and delay parameters using multiple trajectories that reach steady state is accurate and precise when unidentifiable parameters, *K_M_* and *τ_Y_*, are fixed. (**a**) Two-step activation model diagram. The unobserved protein, X, is produced at rate *A_X_*, and activates the production of a downstream protein, Y, after a random delay *τ_X_*. Activation is modeled by a Michaelis-Menten function. After transcription is initiated, protein Y takes a random time, *τ_Y_*, to mature. Both delays, *τ_X_* and *τ_Y_* follow Gamma distributions with distinct parameters. Proteins X and Y are diluted at the same rate, *B*, due to cell growth. We assume that only protein Y is observed, and the protein count is recorded at discrete times. These measurements are corrupted by a combination of additive and multiplicative observational noise, *ε*(*t*), which follows the normal distribution *N*(0, *Y*(*t*) + *σ_e_*). (**b**) 40 noisy trajectories of measurements, *Y*_obs_(*t*), at time 0, 1,…, 100 (min) of the two-step activation model in Fig. 2 with kinetic parameters *A_X_* = 10 min^-1^, *A_Y_* = 60 min^-1^, *K_M_* = 100, B = 0.05 min^-1^, delays *τ_X_* ~ Γ(18/5, 3/5), *τ_Y_* ~ Γ(18/5, 3/5), and observational noise ~ *N*(0, *Y*(*t*) + 10). (**c**) The simulated measurements of *Y*_obs_(*t*) are indistinguishable when varying *A_X_* and *K_M_* while keeping their ratio, *A_X_*/*K_M_*, constant. Here *A_X_*/*K_M_* = 0.1. The upper and lower boundaries of the shaded region correspond to the mean±SD for the 40 trajectories obtained for each parameter set. (**d**) When estimating parameters using the trajectories in (b), we found that the means of the two delays, *μ_τ_X__* and *μ_τ_Y__*, were not individually identifiable, but their sum could be accurately estimated. (**e–h**) To resolve this identifiability issue, we assumed that the distribution of *τ_Y_* could be estimated separately, and the distribution was thus fixed in the estimation process. We could then accurately estimate *A_X_*/*K_M_*, *μ_τ_X__*, *A_Y_*, and *B*. These estimates became more accurate and precise as we increased the number of trajectories used for inference. Here, sample values were divided by true parameter values to obtain posterior distributions of the normalized parameters.

To test our inference algorithm, we generated 40 time series of the observed protein Y (i.e., *Y*(*t*)) using a delayed stochastic simulation algorithm (Cai, 2007), and added combined additive and multiplicative noise, sampled from *N*(0, *Y*(*t*) + 10) at each time *t*, to obtain 40 synthetic measurement trajectories, *Y*_obs_(*t*), that we subsequently used for inference (Fig. 3b). Trajectories were indistinguishable when both parameters *A_X_* and *K_M_* were scaled by the same factor 2 or 0.5 (Fig. 3c). This degeneracy follows directly from the equation defining the production (a) Two-step activation model of *Y*, *A_Y_ X*(*t*)/(*K_M_* + *X*(*t*)): Because the level of *X*(*t*) is proportional to *A_X_*, the production term is proportional to *A_Y_ A_X_*/(*K_M_* + *A_X_*), which can be rewritten as *A_Y_ A_X_*/*K_M_* /(1 + *A_X_*/*K_M_*). Thus, the production of Y is mainly governed by the ratio *A_X_*/*K_M_* rather than the individual parameters *A_X_* and *K_M_*. This degeneracy leads to the unidentifiability of *A_X_* and *K_M_*. We therefore fixed *K_M_* to an arbitrary value, 100, and estimated the ratio *A_X_*/*K_M_* instead of *A_X_* in the following.

When we estimated the remaining parameters, the posterior samples of the mean time delay needed for gene *x* to regulate gene y, *τ_X_*, and the mean synthesis delay of protein Y, *τ_Y_*, were strongly correlated, indicating that their sum could be accurately estimated, but each could not be estimated individually (Fig. 3d). Hence, measurements of Y contain information only about the sum of the two mean time delays. Indeed, simulated trajectories of the model with *τ_X_* ~ Γ(3.6, 0.6) and *τ_Y_* ~ Γ(3.6, 0.6), both with means equal to 6 mins, are indistinguishable from simulations with *τ_X_* ~ Γ(1.2, 0.6) and *τ_Y_* ~ Γ(6.0, 0.6), with means of 2 and 10 mins, respectively (Supplementary Fig. **S1**). This is consistent with the fact that the total delay in a cascade equals the sum of individual delays in the deterministic case (Glass *et al*., 2021).

The expression delay of Y, *τ_Y_*, can be directly estimated using an independent experiment with a genetic circuit containing only gene *Y*. We therefore assumed that the distribution of *τ_Y_* is known to resolve the identifiability issue with *τ_X_* and *τ_Y_*. As a result, we obtained accurate and precise estimates of all remaining parameters: *A_X_*/*K_M_*, *A_Y_*, *τ_X_*, and *B* (Fig. 3e–h). These estimates became more precise when we increased the number of measured trajectories. Precision also increased with measurement frequency (Supplementary Fig. **S2**).

Thus tests with a simple circuit and synthetic data suggest that it is in general impossible to infer all parameters in biochemical reaction networks when some of the components are not observed. Some of the unidentifiable parameters need to be fixed to accurately estimate the remaining parameters in the two-step activation model. For synthetic gene circuits, some of these parameters could be estimated in separate experiments, while only combinations of other parameters may be inferable. This identifiability analysis was possible because we used a Bayesian approach and estimated the joint posterior of the parameter (Hines *et al*., 2014).

### 3.2 With short time series, to estimate kinetic and delay parameters, the decay rate needs to be known

In time-lapse microscopy experiments cells can enter stationary phase (Kolter *et al*., 1993), or the experiment may last too long before the population reaches an equilibrium distribution. To test whether the kinetic and delay parameters can be accurately estimated from data obtained over shorter time intervals we applied our algorithm to synthetic data corresponding to the first half of the experiment we described in the previous sections (Fig. 4a).

**Fig. 4.**
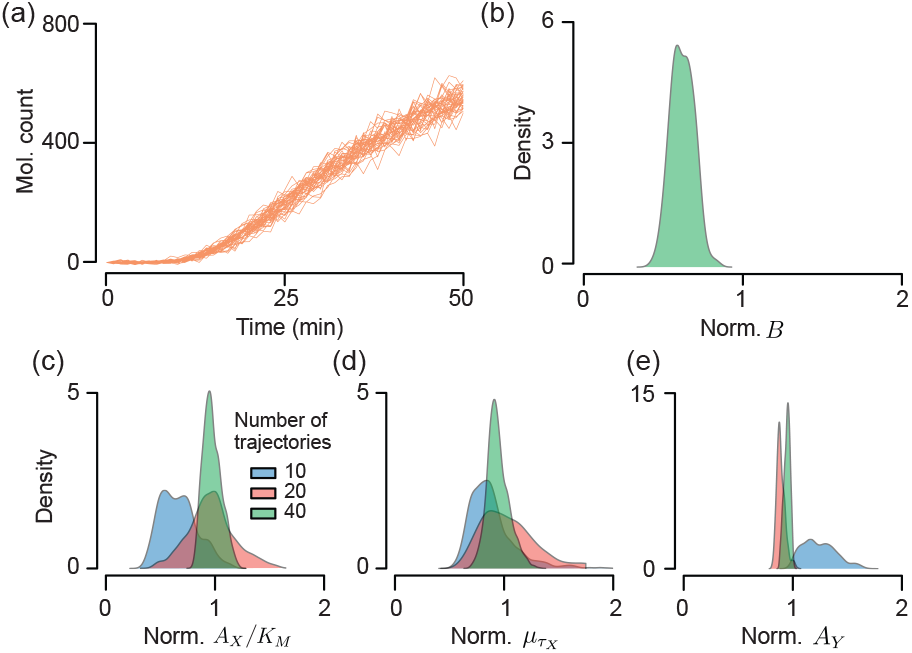
Estimation of the kinetic and delay parameters using multiple short trajectories that did not reach steady state becomes accurate when the decay rate, *B*, is assumed known. (**a**) 40 trajectories of noisy measurements at times 0, 1,…, 50 (min) of the two-step activation model shown in Fig. 2 using the same parameter values as in Fig. 3a. Here, the observation window is too short for the simulated trajectories to reach steady state, unlike those in Fig. 3a. (**b**) We also assumed that K_M_ and the distribution of *τ_Y_* are known, and found that the dilution rate, *B*, is underestimated using our algorithm. This indicates that the decay rate, *B*, can be accurately estimated only when measurements are taken until the system reaches steady state. (**c–e**) To account for this bias, we assume that the decay rate, *B*, is known, and fixed it at its true value in our inference algorithm. This allowed us to obtain the accurate and precise estimates of the ratio *A_X_*/*K_M_*, μ_τ_X__, and *A_Y_*, which improved with the number of measurement trajectories. Sample values were divided by true parameter values to obtain posterior distributions of the normalized parameters.

Under the conditions leading to accurate and precise estimation in the previous section (i.e., assuming that *K_M_* and the distribution of *τ_Y_* are known), we observed that the posterior mean of the decay rate, *B*, was considerably lower than the true parameter value (Fig. 4b). This underestimate occurred because the decay rate does not strongly affect the dynamics of the observed and unobserved proteins in the transient regime before their counts reach steady state values, since the terms *BX*(*t*) and *BY*(*t*) are small until protein counts increase. Thus, B needs to be measured separately to estimate the other parameters accurately. If the decay mainly occurs via growth-induced dilution, B can be estimated by measuring single-cell growth trajectories obtained with time-lapse microscope. By fixing the decay rate to its true value, we obtained accurate estimates for the other parameters *A_X_*/*K_M_*, μ_τ_X__, and *A_Y_*, and these became more precise as we increased the number of measurement trajectories (Fig. 4c–e).

We therefore found that the dilution rate needs to be estimated separately to obtain accurate and precise estimates of the remaining kinetic and delay parameters in the two-step activation model (Fig. 3a) when observed trajectories did not reach a steady state. We expect that similar identifiability issues will persist in more complex models.

### 3.3 Estimation of time delays in unobservable transcriptional regulation

We applied our method to data obtained from a two-step activation circuit in *E. coli* (Fig. 5a). The time-lapse fluorescence images of the cell populations were obtained previously (Cheng *et al*., 2017). The two-step activation circuit consists of two genes; one encodes the unobserved transcriptional activator AraC and the second encodes the observed YFP, corresponding to X and Y in the two-step activation model, respectively (Fig. 3a). Once IPTG and arabinose are added at time *t* = 0, AraC is activated. After maturation of the expressed AraC, the mature protein searches a downstream target binding site and initiates the synthesis of YFP. The single-cell fluorescence signal from matured YFP was measured over 50 min (Fig. 5b). From the measured fluorescence intensity for two independent experiments with 23 and 25 cells, we obtained molecular counts by multiplying a previously calculated conversion rate (Fig. 5c) (Choi *et al*., 2020). Because these trajectories did not reach a steady state, we first estimated the decay rate, *B*, and the time delay for the synthesis of YFP, *τ_Y_*, to be able to accurately estimate the remaining kinetic and delay parameters from data (Fig. 3d and 4b).

**Fig. 5.**
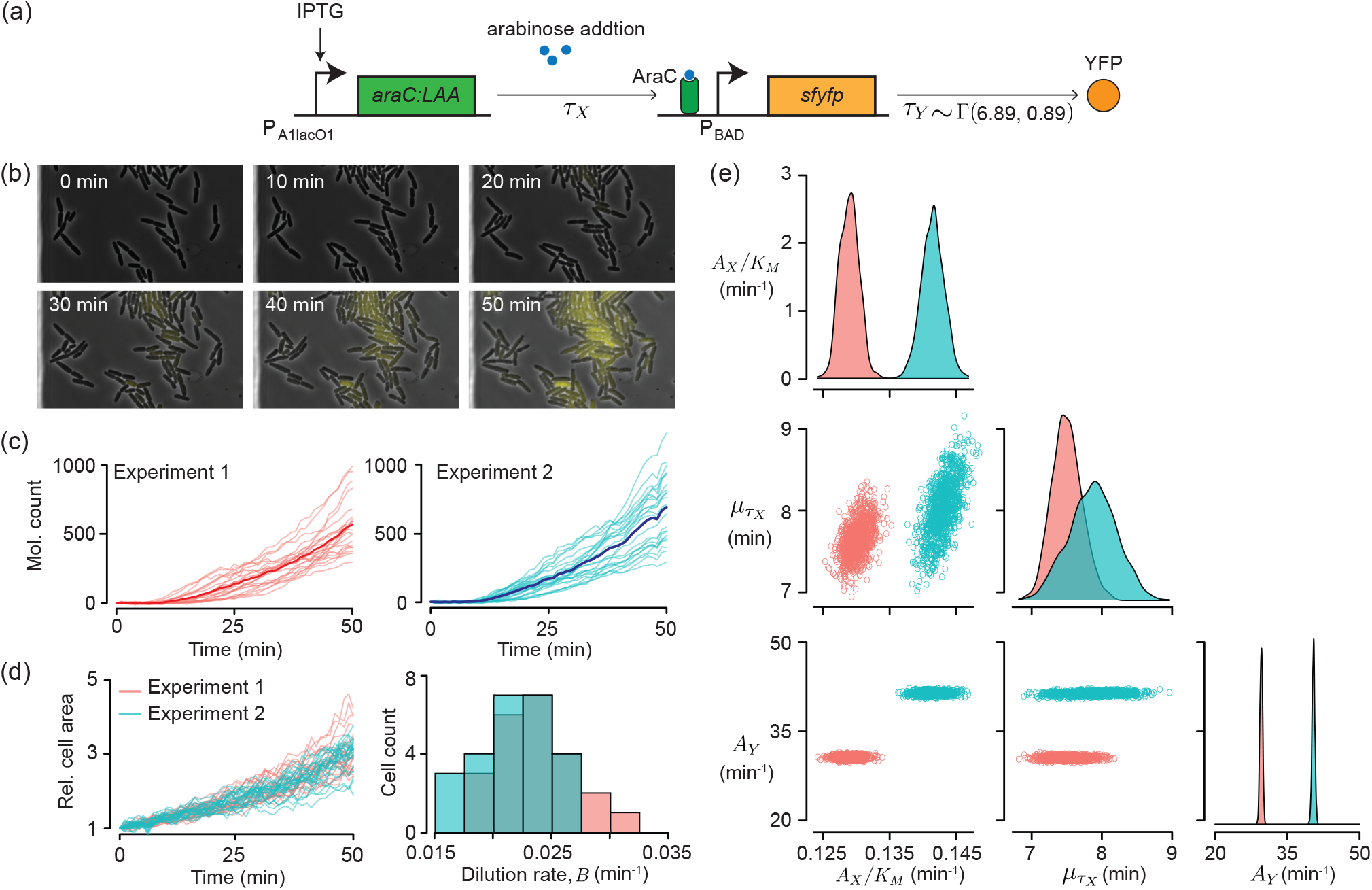
Our inference method provided similar estimates of the mean regulation delay from two independent experiments with a two-step activation circuit in *E. coli*. (**a**) The two-step activation circuit in E. *coli*. Once IPTG and arabinose are added to the growth media, the expression of *araC* is induced and it activates the synthesis of YFP after the time delay of the regulation, *τ_X_*. The synthesis of YPF also involves a time delay, *τ_Y_* ~ Γ(6.89, 0.89), which we estimated previously using a reporter-only circuit. In this circuit, only the fluorescence level of YFP is measured while the level of AraC is not measurable. (**b**) Time-lapse images of YFP expression from the two-step activation circuit monitored using fluorescence microscopy (Cheng *et al*., 2017). The fluorescent cells were observed after induction with 0.2mM IPTG and 2% (w/v) arabinose at time 0. (**c**) Molecular counts were obtained by dividing the fluorescence level of each cell by a conversion constant, calculated in our previous paper (Choi *et al*., 2020). The numbers of cell trajectories are 23 and 25, respectively. The thick lines represent the mean trajectories. (**d**) The area of each cell from two independent experiments was measured by tracking the lineage of each cell. When a mother cell divided into two daughter cells, the area of the mother cell was added to the areas of its daughter cells. Additionally, the area was normalized by its initial value, obtaining the relative cell area of each cell (left). The relative areas were used to estimate the dilution rate, B, by fitting an exponential function to the relative area trajectories (right). (**e**) Applying our inference method to the cell trajectories, we obtained the posterior samples of the parameters *A_X_*/*K_m_*, *μ_τ_X__*, and *A_Y_*. To avoid bias and identifiability issues in estimation, we fixed the dilution rate *B* to its average value, 0.022, as it was directly estimated from the observed cell areas (d) and the time delay of the synthesis of YFP *τ_Y_* to Γ(6.89, 0.89) as it was previously estimated (Choi *et al*., 2020). We obtained the estimates (mean±SD) of *A_X_*/*K_M_*, 12.90±0.14 min^-1^ and 14.15±0.17 min^-1^, and the estimates of *A_Y_*, 30.64±0.27 min^-1^ and 41.45±0.28 min^-1^. These estimates are higher in the second experiment because the mean trajectory is higher in the second experiment (c). On the other hand, the estimates of the mean delay, 7.50±0.21 min and 7.87±0.34 min, were similar in both experiments.

The time delay for the synthesis of YFP can be separately estimated using a ‘reporter-only’ circuit which consists of a *P*_bad_ promoter that drives the expression of YFP without the need for accumulation of AraC. Previously, we used data from experiments with such a circuit to obtain the estimate τ_Y_ ~ Γ(6.89, 0.89) (Choi *et al*., 2020). We used this estimate in the following. For the decay rate, we can use the dilution rate because dilution is the main driver of decay since YFP is stable and is not enzymatically degraded (Andersen *et al*., 1998). The dilution rate can be directly estimated from cell area tracked with a time-lapse microscope (Fig. 5d left) (Megerle *et al*., 2008; Taheri-Araghi *et al*., 2015). We fit an exponential function to each single-cell growth trajectory to estimate individual dilution rates (Fig. 5d right). We used the average of these estimates, 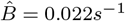, as the dilution rate for the follwing estimates.

To estimate the remaining parameters we applied our MCMC algorithm to the measurements of the observed fluorescent protein and obtained estimates of the posterior distributions for the parameters *A_X_*/*K_M_*, *μ_τ_X__*, and *A_Y_*. The posterior means of the kinetic parameter *A_X_*/*K_M_* and *A_Y_* in Experiment 1 were higher than those in Experiment 2 (Fig. 5e). This difference was due to a ~20% higher intensity in Experiment 1 compared to Experiment 2 (Fig. 5c). On the other hand, the posterior means of the expected time delay for the transcriptional regulation of AraC,, are similar between the two experiments: 7.50±0.21 min and 7.87±0.34 min (Fig. 5e).

The estimated time delay for the transcriptional regulation is the sum of delays corresponding to AraC synthesis, diffusion, and binding-site search. Interestingly, the binding-site search time of transcriptional factors in prokaryotic cells is usually less than a few minutes even for chromosomal genes whose copy number is just one or two (Hammar *et al*., 2012; Elf *et al*., 2007). The two-step activation circuit is plasmid-borne, and thus has copy number in the dozens. This indicates that the binding-site search time is much shorter than those for chromosomal genes (i.e., less than a minute), so the estimated time delay (~7.5 min.) mostly comes from the synthesis, including transcription, translation, folding, and dimerization. This time delay has not been characterized previously. This finding thus provides a better understanding of the kinetics of AraC protein, a widely used transcriptional activator in synthetic biology.

## 4 Conclusion

We have developed a simulation-based Bayesian MCMC method for the inference of kinetic and delay parameters from noisy measurement of a GRN with unobserved components. We applied the method to a two-step activation model where an unobserved species X regulates the synthesis of an observed species Y.

Using synthetic data, we have shown that certain parameters are not identifiable. However, accurate estimates of some of these parameters can be obtained if the *observable* system components can be characterized in separate experiments. Specifically, we found that the production rate of X, A_X_, and the Michaelis-Menten constant for regulating gene Y, K_M_, cannot be estimated independently. However, the ratio between these two parameters (i.e., *A_X_*/*K_M_*) is identifiable from the observed trajectories (Fig. 3c). We also found that the sum of two delays, the regulatory delay, *τ_X_*, and synthesis delay, *τ_Y_*, was identifiable, but the delays were not identifiable individually (Fig. 3d). Often *τ_Y_* or *K_M_* can be estimated from separate experiments. In that case the regulatory delay, *τ_X_*, and production rate, *A_X_*, are identifiable with data from the full circuit. Furthermore, if the measured trajectories do not reach steady state, the decay rate, B, is also not identifiable (Fig. 4b). For proteins that are not actively degraded this last identifiability issue can be resolved by estimating the decay rate directly from cell size measurements.

We applied our method to two independent sets of experimental timelapse fluorescence data from a two-step activation circuit in which AraC protein activates the synthesis of YFP. We fixed the decay rate to the dilution rate estimated from the observed cell area time series (Fig. 5d). Furthermore, we also fixed the delay for the synthesis of YFP to the value estimated with YFP-reporter only circuit in our previous work (Choi *et al*., 2020). This allowed us to estimate the production rates and regulation delay parameters of unobserved AraC protein using only observations of YFP. The estimated production rates *A_X_*/*K_M_* and *A_Y_* were higher in the second experiment (Fig. 5e). This might be due to the mean trajectory being higher in the second group (Fig. 5c). On the other hand, the estimated regulation delays, *τ_X_*, are similar in the two experiments. This might be because regulation delay is an inherent attribute of AraC and the downstream gene *sfyfp* unlike the fluorescence level, which can be sensitive to camera settings. We hypothesize that the estimated time delay of transcriptional regulation of AraC (~ 7.5 min.) mostly comes from the synthesis, not binding-site search. This finding can play an important role in building synthetic circuits as the AraC protein, whose kinetics are yet to be fully characterized, is a widely used transcriptional activator in synthetic biology (Romano *et al*., 2021; Moon *et al*., 2012).

For example, transcriptional regulators are being used in constructing cascaded genetic logic gates or oscillators where delay can impact output and dynamics (Moon *et al*., 2012; Mather *et al*., 2014).

As the number of parameters and unobserved species in a system increases, identifiability issues often worsen (Raue *et al*., 2009; Hines *et al*., 2014; Browning *et al*., 2020). In addition, more complex models typically result in more local maxima in the corresponding posterior distributions. Markov chains can get trapped at a local maximum, leading to inaccurate estimates. This problem could be resolved by adopting advanced sampling methods, such as the multiset samplers (Leman *et al*., 2009; Kim and MacEachern, 2015) which relies on many agents in one Markov chain and can more easily escape a local maximum.

While our method may appear similar to approximate-Bayesian computation (ABC) approach (Beaumont *et al*., 2002), the two are qualitatively different. While in our method the proposed reaction counts are accepted based on the MH acceptance probability, in the ABC approach a proposed sample is accepted based on a metric and a threshold, both of which need to be defined appropriately. Our method is related to the ABC-MCMC approach (Marjoram *et al*., 2003), in which parameters are accepted stochastically. However, ABC-MCMC uses stochastic simulations only for computing the acceptance probability to update a proposed parameter while in our approach a stochastic simulation of the model system is used to obtain proposed reaction counts.

Thus, the key step of the present method is the use of a stochastic simulation algorithm. The acceptance probability of proposed reaction counts based on the MH algorithm can be computed using the likelihood ratio of only observed processes (Eq. 10), independently of the complex dynamical model of the system. This allows for our method to be easily implemented. Furthermore, this way to obtain proposal reaction counts captures strong correlations between individual counts without the need to tune hyperparameters. In contrast, a direct application of the block-update method (Boys *et al*., 2008; Choi *et al*., 2020; Cortez *et al*., 2022) requires the tuning of numerous proposal distributions, leading to small acceptance rates and slow convergence of the MCMC.

We assumed that all cells in a population are identical, and thus share the same parameters, i.e., we ignored the cell-to-cell variability. Even isogenic cell populations can show significant cell-to-cell variability (Kepler and Elston, 2001; Kærn *et al*., 2005; Raj and van Oudenaarden, 2008; Smith and Grima, 2018), and heterogeneity in a population plays a crucial role in biological processes such as development. Our method can be extended to account for such variability using hierarchical model, a potential avenue for future work.

Our approach is scalable. Because the likelihood function has been derived for a general biochemical reaction network, the simulation-based MCMC can be tailored to a more complex model. The method does require performing stochastic simulations of the biochemical system for each MCMC iteration, resulting in a high computational cost. This cost could be reduced by developing an emulator, which is a fast data generator replacing a slower computational model to avoid the sampling process (Kennedy and O’Hagan, 2001).

A simulation-based MCMC method can be developed for other dynamical models as long as a likelihood function is available. While we used a continuous-time Markov Chain, which efficiently explains a biochemical reaction network with a low copy number of molecules, one can also use a stochastic differential equation (Calderazzo *et al*., 2018; Ruttor and Opper, 2009) which is accurate when the copy numbers are higher, an agent-based model (Grazzini et al., 2017), or a delay differential equation (Kim *et al*., 2022). Thus, we expect that our framework can be extended to various stochastic models of non-Markovian GRNs, and thus characterize the dynamics of a variety of systems from partial observations.

## Data Availability

All experimental data has been published previously, and is available upon request.

